# A Mammalian High-Throughput Screen for AI-Designed Peptide-Guided Protein Degraders

**DOI:** 10.64898/2026.08.24.746873

**Authors:** Lin Zhao, Anuoluwapo Mattix, Aastha Pal, Tong Chen, Sophia Vincoff, Lauren Hong, Diana Renteria, Sunetra Sase, Adeline L. Vanderver, Daniel R. Matson, Pranam Chatterjee

## Abstract

Targeted protein degradation (TPD) offers a route to eliminate disease-driving proteins that remain inaccessible to conventional inhibitors. However, degrader discovery remains low-throughput, labor-intensive, and dependent on randomized libraries or non-human display systems, limiting functional selection in mammalian cells. Here, we present a high-throughput, human cell-based platform for screening peptide-guided ubiquibodies (uAbs). These genetically encodable, doxycycline-inducible degraders fuse peptide guides generated by protein language models to the CHIPΔTPR E3 ligase domain, creating a modular, CRISPR-like system for programmable TPD. For each target, we introduce a pooled uAb library into the corresponding fluorescent reporter cell line, isolate cells with reduced target abundance by FACS, and recover enriched peptide guides by sequencing. For *β*-catenin, enriched uAbs reduced endogenous *β*-catenin abundance and Wnt signaling in DLD1 cells. GFAP-directed uAbs reduced endogenous GFAP abundance and cell viability in U251 glioblastoma cells, while EWS::FLI1-directed uAbs reduced fusion oncoprotein abundance, suppressed EWSAT1 expression, and increased apoptosis in Ewing sarcoma models. Finally, a screen using endogenously tagged GATA2 further identified uAbs that reduced GATA2 under native genomic regulation. Overall, our platform connects generative peptide design to functional mammalian selection and establishes a scalable strategy for CRISPR-like proteome perturbation.

## Introduction

A substantial fraction of disease-driving proteins, including transcription factors and fusion oncoproteins, remain inaccessible to small molecules because disordered regions and poorly defined surfaces lack conventional binding pockets [1, 2]. Antibodies, nanobodies, and peptides can recognize these surfaces, and display technologies have enabled their discovery from large randomized libraries [2, 3, 4, 5, 6]. These workflows remain labor-intensive and select binders outside the mammalian intracellular context in which function must ultimately be established.

Structure-based generative models have accelerated binder design for targets with defined conformations [7, 8, 9, 10]. Their reliance on specified three-dimensional structures, however, complicates design against disordered proteins and selective recognition of related targets or conformational states. Sequence-based protein language models (pLMs), including PepMLM, PepPrCLIP, and SaLT&PepPr, address this limitation by designing peptide binders directly from target sequences, including for conformationally diverse proteins [11, 12, 13]. More recent generative and guidance frameworks, including moPPIt, SOAPIA, and AlloGen, enable motif-, isoform-, and conformation-specific design [14, 15, 16], while sequence-based diffusion and flow-matching frameworks have further supported multi-objective optimization of binding and developability [17, 18, 19, 20].

Beyond occupancy-based target inhibition, peptide guides generated by these methods have been fused to the CHIPΔTPR E3 ligase domain to create ubiquibodies (uAbs) that direct target ubiquitination and proteasomal degradation [21, 22, 23]. In this CRISPR-like proteome-editing architecture, the peptide serves as the targeting guide and the E3 ligase domain as the effector. However, rapidly identifying peptide guides that produce functional degradation in mammalian cells remains a major bottleneck.

High-throughput CRISPR screens achieve scale by coupling sequence-defined guide RNA libraries to selectable cellular phenotypes and recovering enriched or depleted guides by sequencing [24, 25, 26]. Here, we apply this logic to protein degradation by establishing a pooled mammalian screen for functional uAbs. For each target, we introduce a separate doxycycline-inducible uAb library into the corresponding fluorescent reporter cell line, isolate cells with reduced target fluorescence by FACS, and identify enriched peptide guides via next-generation sequencing for downstream functional validation.

We selected four targets to evaluate distinct capabilities of the platform. We first used *β*-catenin, for which our prior studies had generated functional peptide guides, to establish the screen [12, 13, 27]. Enriched uAbs reduced endogenous *β*-catenin and suppressed Wnt signaling, with proteasome-dependent degradation that increased over 72 h. We then tested GFAP and EWS::FLI1 as disease-relevant targets that remain difficult to address through conventional ligand discovery. GFAP accumulation causes Alexander disease, while the extensively disordered EWS::FLI1 fusion oncoprotein drives Ewing sarcoma and presents a stringent test of sequence-based peptide design [28, 29, 30, 31]. GFAP-directed uAbs reduced endogenous GFAP and cell viability in U251 glioblastoma cells, whereas EWS::FLI1-directed uAbs reduced EWS::FLI1 abundance, suppressed EWSAT1 expression, and increased apoptosis in Ewing sarcoma cells. Finally, candidate uAbs reduced mNeonGreen-tagged GATA2 expressed from its native genomic locus [32, 33]. Together, these studies establish a scalable route from generative peptide design to CRISPR-like functional proteome perturbation.

## Results

### A dual-reporter system enables pooled screening of peptide-guided degraders

To select peptide guides according to intracellular degradation activity, we engineered a dual-reporter system that couples target abundance to mCherry fluorescence and uAb expression to GFP fluorescence. We first introduced lentiviral constructs encoding target-mCherry fusions and selected monoclonal reporter lines with stable expression. We then transduced each reporter line with a doxycycline-inducible uAb library in which a model-designed peptide guide was fused to the CHIPΔTPR E3 ligase domain (Figure 1A). Fluorescence microscopy confirmed stable mCherry expression in the *β*-catenin, GFAP, and EWS::FLI1 reporter lines (Figure 1B) and doxycycline-dependent induction of the uAb-GFP cassette (Figure 1C).

**Figure 1:**
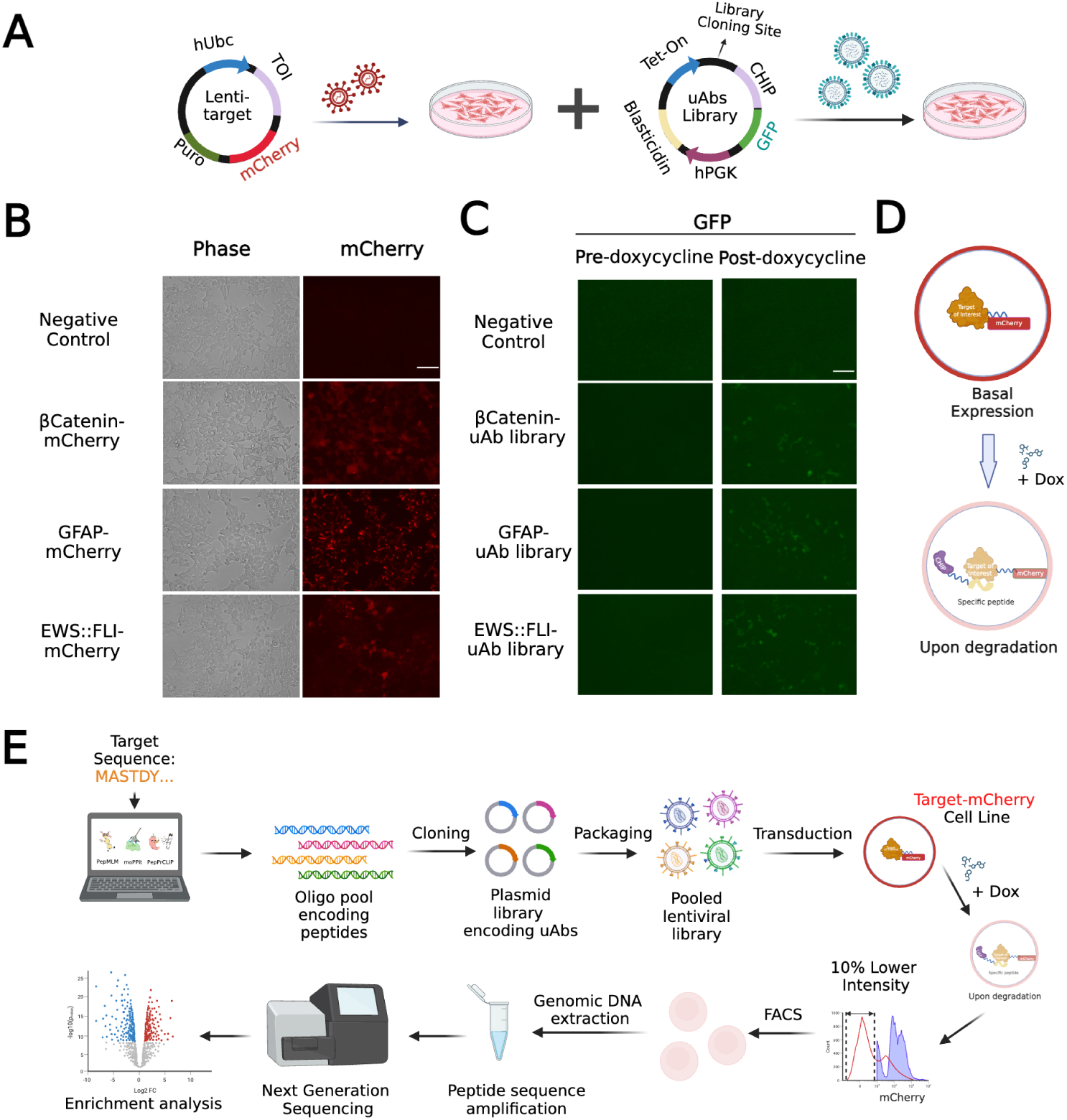
A dual-reporter system for pooled mammalian screening of peptide-guided uAbs. **(A)** Lentiviral delivery of the target-mCherry reporter and doxycycline-inducible uAb-GFP library constructs. **(B)** Representative phase-contrast and mCherry fluorescence images of the negative-control, *β*-catenin-mCherry, GFAP-mCherry, and EWS::FLI1-mCherry reporter lines. **(C)** GFP fluorescence before and after doxycycline treatment, confirming induction of the corresponding uAb libraries. **(D)** Schematic of peptide-guided recruitment of CHIPΔTPR to a target-mCherry fusion, resulting in target degradation and reduced mCherry fluorescence. **(E)** Pooled screening workflow comprising computational peptide design, oligonucleotide synthesis, uAb-library cloning, lentiviral packaging, reporter-cell transduction, doxycycline induction, FACS isolation of reporter-low cells, recovery of peptide-encoding sequences, next-generation sequencing, and enrichment analysis. Scale bars in **B** and **C**, 125 *µ*m.

In this system, a functional peptide guide recruits the CHIPΔTPR domain to the target-mCherry fusion, leading to target ubiquitination, degradation, and reduced mCherry fluorescence (Figure 1D). For each target, we synthesized a library of 2,000 20-amino-acid guides designed using PepMLM, PepPrCLIP, or moPPIt [11, 12, 14] (Table S1). We cloned each oligonucleotide pool into the uAb vector, packaged the resulting plasmid library into lentivirus, transduced the corresponding reporter line, and induced uAb expression with doxycycline. We then isolated cells with the lowest 10% of target-reporter fluorescence by FACS and quantified peptide-encoding sequences by next-generation sequencing (Figure 1E). The gating strategy selected singlet, reporter-positive cells before collection of the target-reporter-low population (Supplementary Figure S1). Thus, the dual-reporter system converted intracellular degradation into a selectable fluorescence phenotype linked to each peptide sequence.

### *β*-catenin establishes screen performance and functional degradation

We first evaluated our platform using *β*-catenin as a target, as prior studies have generated functional peptide-guided degraders against this target, including *β*cat_SnP_8 uAb [23, 12]. This established positive control allowed us to determine whether fluorescence enrichment recovered a known degrader while identifying additional functional guides. We compared peptide counts between pre-sort and reporter-low populations using MAGeCK and ranked each guide by its log_2_ fold change and associated *p*-value [34, 35]. The resulting analysis recovered *β*cat_SnP_8 and identified additional significantly enriched guides (Figure 2A).

**Figure 2:**
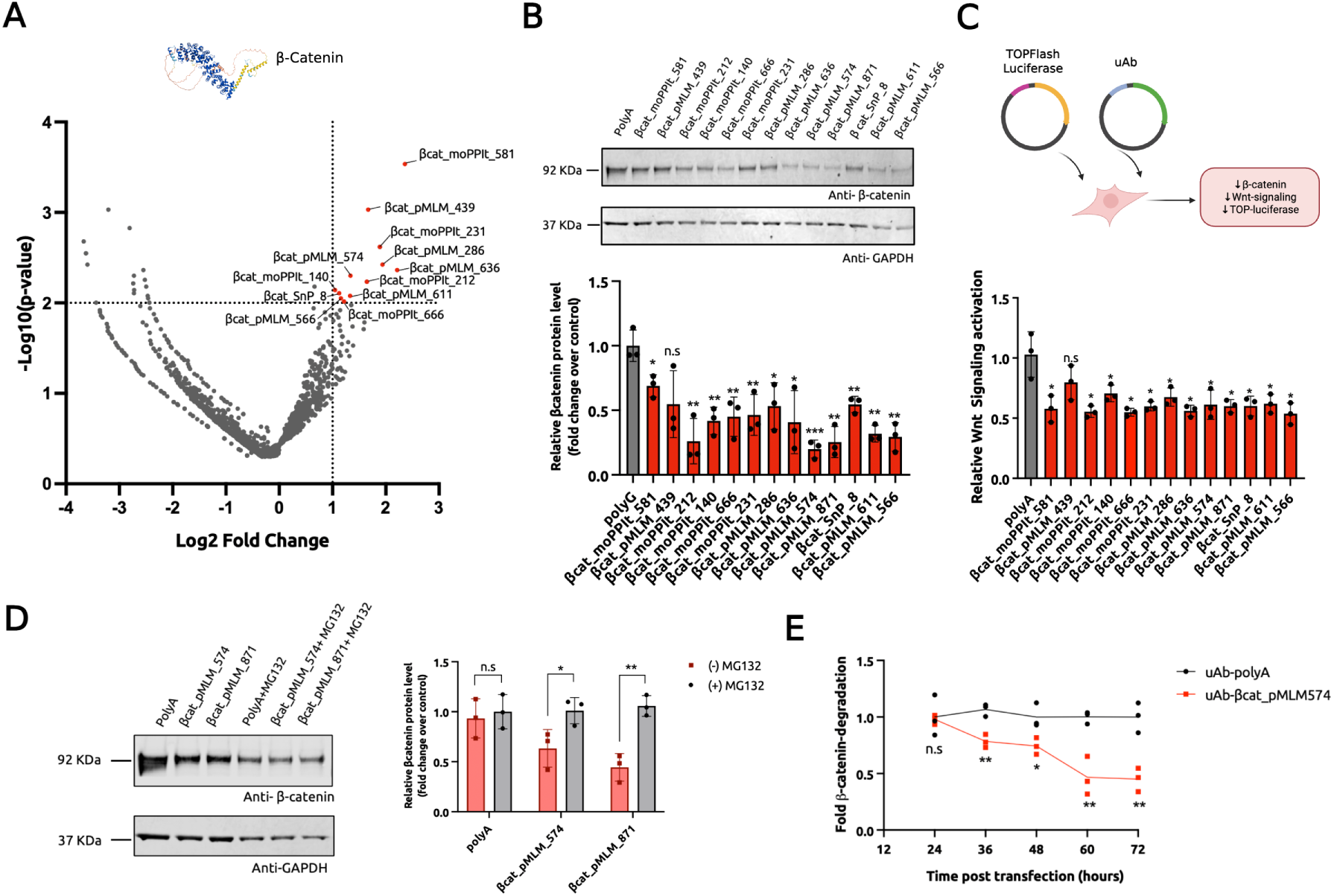
Pooled screening and functional validation of *β*-catenin-directed uAbs. **(A)** Volcano plot of peptide enrichment in the *β*-catenin-mCherry reporter screen. Red points indicate significantly enriched peptides with *p <* 0.01 and log_2_ fold change *>* 1. **(B)** Immunoblotting and densitometric quantification of endogenous *β*-catenin in DLD1 cells expressing selected enriched uAbs, the previously validated *β*cat_SnP_8 positive control, or the polyA negative control. **(C)** TOPFlash luciferase measurements of Wnt-dependent transcription in DLD1 cells expressing the indicated uAbs. **(D)** Immunoblotting and densitometric quantification of endogenous *β*-catenin in DLD1 cells expressing *β*cat_pMLM_574 or *β*cat_pMLM_871 in the presence or absence of MG132. **(E)** Time-dependent change in endogenous *β*-catenin abundance in DLD1 cells expressing *β*cat_pMLM_574 or polyA. Cells were collected at 24, 36, 48, 60, and 72 h after transfection. Bars and points report mean *±* s.d. from three independent experiments (*n* = 3).

To determine whether reporter enrichment predicted degradation of the untagged endogenous protein, we reconstructed selected uAbs and expressed them in DLD1 colorectal cancer cells. Immunoblotting showed that several enriched uAbs reduced endogenous *β*-catenin relative to the non-targeting polyA control (Figure 2B). We then measured Wnt-dependent transcription using a TOPFlash luciferase reporter [36]. Multiple uAbs reduced TOPFlash activity, demonstrating that target loss produced the expected functional suppression of Wnt signaling (Figure 2C).

We next tested whether the observed target loss required proteasomal activity. MG132 attenuated the reductions produced by *β*cat_pMLM_574 and *β*cat_pMLM_871, supporting degradation through the ubiquitin-proteasome pathway (Figure 2D). A 72-h time course further showed a progressive reduction in endogenous *β*-catenin following expression of *β*cat_pMLM_574 (Figure 2E).

To assess whether enrichment distinguished active from inactive guides, we selected five non-enriched peptides from the *β*-catenin screen (Supplementary Figure S2A). None reduced endogenous *β*-catenin abundance (Supplementary Figure S2B) or significantly changed TOPFlash activity (Supplementary Figure S2C). Time-course immunoblotting independently confirmed the progressive activity of the enriched *β*cat_pMLM_574 uAb relative to polyA (Supplementary Figure S2D). Together, these experiments established that the screen recovered known and newly designed *β*-catenin degraders whose activity extended from reporter selection to endogenous degradation, pathway suppression, proteasome dependence, and defined degradation kinetics.

### GFAP-directed uAbs reduce target abundance and U251 cell viability

Having established the screen with *β*-catenin, we next tested whether it could identify degraders against GFAP, a disease-associated intermediate filament protein for which peptide-guided degradation had not been established. Pathogenic GFAP accumulation drives Alexander disease, and its filamentous architecture presents a distinct targeting challenge [28, 29]. Enrichment analysis identified multiple candidate guides in the GFAP-mCherry reporter screen (Figure 3A). We reconstructed selected enriched uAbs and expressed them in U251 glioblastoma cells, which endogenously express GFAP. Immunoblotting showed that multiple candidates reduced endogenous GFAP abundance relative to the control uAb (Figure 3B).

**Figure 3:**
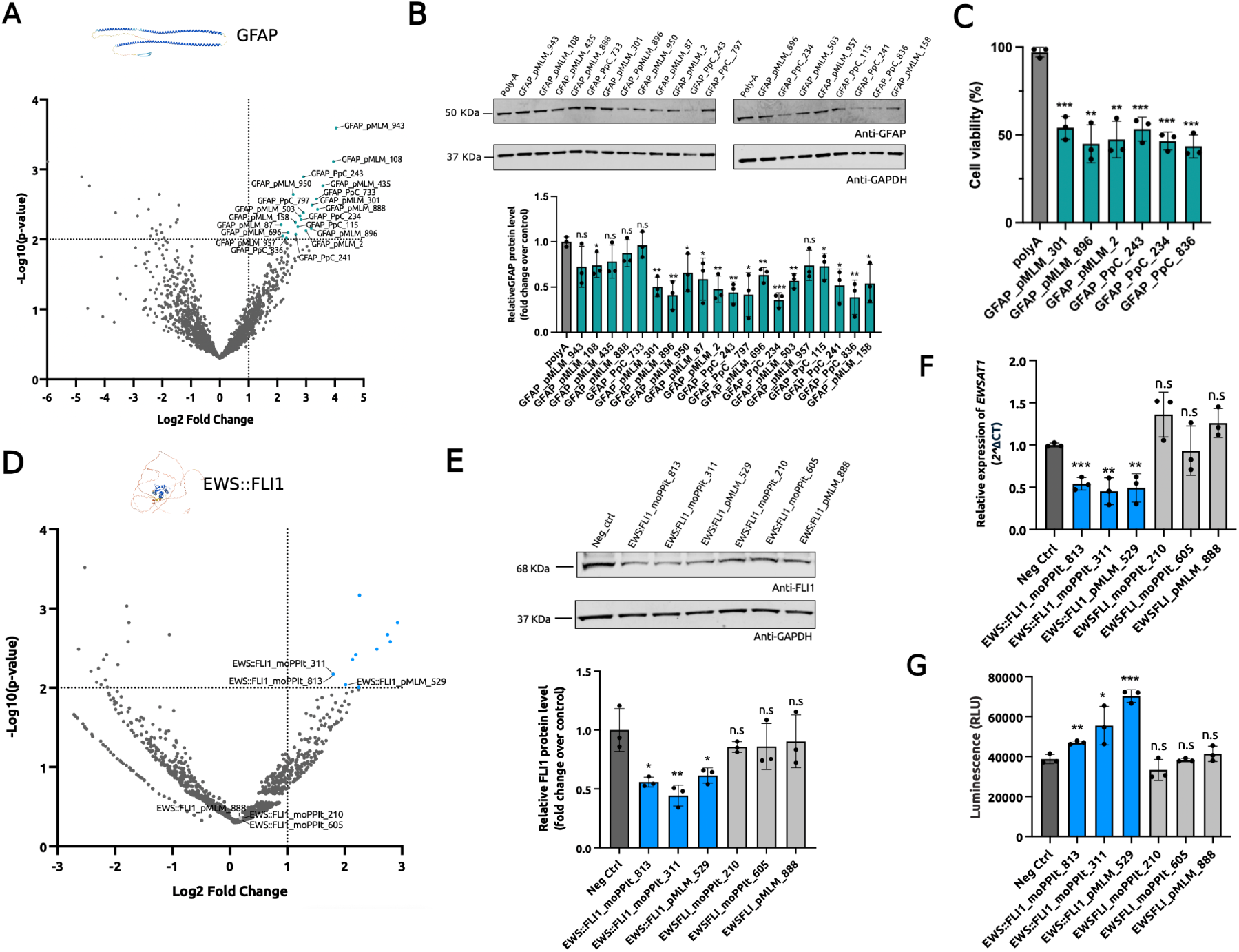
Screening and functional validation of GFAP- and EWS::FLI1-directed uAbs. **(A)** Volcano plot of peptide enrichment in the GFAP-mCherry reporter screen. Cyan points indicate significantly enriched peptides with *p <* 0.01 and log_2_ fold change *>* 1. **(B)** Immunoblotting and densitometric quantification of endogenous GFAP in U251 cells expressing selected enriched GFAP-directed uAbs. **(C)** Cell viability of U251 cells following expression of six selected GFAP-directed uAbs or the polyA control. **(D)** Volcano plot of peptide enrichment in the EWS::FLI1-mCherry reporter screen. Orange points indicate significantly enriched peptides with *p <* 0.01 and log_2_ fold change *>* 1. **(E)** Immunoblotting and densitometric quantification of endogenous EWS::FLI1 in TC71 cells (via an anti-FLI1 antibody) expressing three enriched or three non-enriched EWS::FLI1-directed uAbs. **(F)** Relative EWSAT1 expression measured by quantitative RT-PCR in TC71 cells expressing the indicated uAbs. **(G)** Annexin V apoptosis measurements in TC71 cells expressing the indicated uAbs. Blue bars in **E-G** denote enriched candidates, and gray bars denote non-enriched candidates. Bars report mean *±* s.d. from three independent experiments (*n* = 3).

We then asked whether GFAP degradation altered U251 cell fitness. Six selected GFAP-directed uAbs reduced cell viability relative to polyA, connecting endogenous target loss to a cellular phenotype (Figure 3C). As an orthogonal test of screen discrimination, we selected five non-enriched guides from the GFAP screen (Supplementary Figure S3A). None significantly reduced endogenous GFAP in U251 cells (Supplementary Figure S3B). These results extended the screen beyond a previously validated target and identified functional uAbs against a disease-relevant filament protein.

### Sequence-designed uAbs perturb the disordered EWS::FLI1 oncoprotein

We then sought to determine whether sequence-based peptide design and mammalian selection could reach an extensively disordered fusion oncoprotein. EWS::FLI1 is the principal oncogenic driver of Ewing sarcoma and lacks the stable binding pockets required by many conventional ligand-discovery methods [31, 37, 30]. Its conformational disorder therefore provided a stringent test of the sequence-based design strategy [12, 30].

The EWS::FLI1-mCherry screen identified sixteen enriched guides, including candidates generated by PepMLM and moPPIt (Figure 3D). We selected three enriched and three non-enriched guides and expressed the corresponding uAbs in TC71 Ewing sarcoma cells [38]. All three enriched uAbs reduced endogenous EWS::FLI1 abundance, whereas the three non-enriched candidates did not produce significant degradation (Figure 3E).

To determine whether degradation suppressed EWS::FLI1 function, we measured EWSAT1, a canonical transcriptional target of the fusion oncoprotein [39]. Each enriched uAb reduced EWSAT1 expression, whereas the non-enriched candidates did not (Figure 3F). Annexin V measurements further showed increased apoptosis following expression of the enriched uAbs, with no significant increase for the non-enriched candidates (Figure 3G). Thus, sequence-designed peptide guides supported degradation and functional perturbation of a highly disordered fusion oncoprotein in a disease-relevant cellular model.

### Endogenous GATA2 tagging enables screening under native genomic regulation

The target-mCherry reporters provided a strong screening signal but required ectopic target expression. We therefore asked whether uAb screening could identify degraders against a target expressed from its native genomic locus. Using CRISPR/Cas9-mediated homology-directed repair, we inserted mNeonGreen at the C terminus of endogenous GATA2 in Ark1 uterine serous carcinoma cells (Figure 4A). Immunoblotting with anti-GATA2 and anti-mNeonGreen antibodies confirmed expression of the higher-molecular-weight GATA2-mNeonGreen fusion in the selected clones while retaining untagged GATA2 from the second allele (Figure 4B). Fluorescence microscopy further confirmed nuclear localization of the tagged protein (Figure 4C).

**Figure 4:**
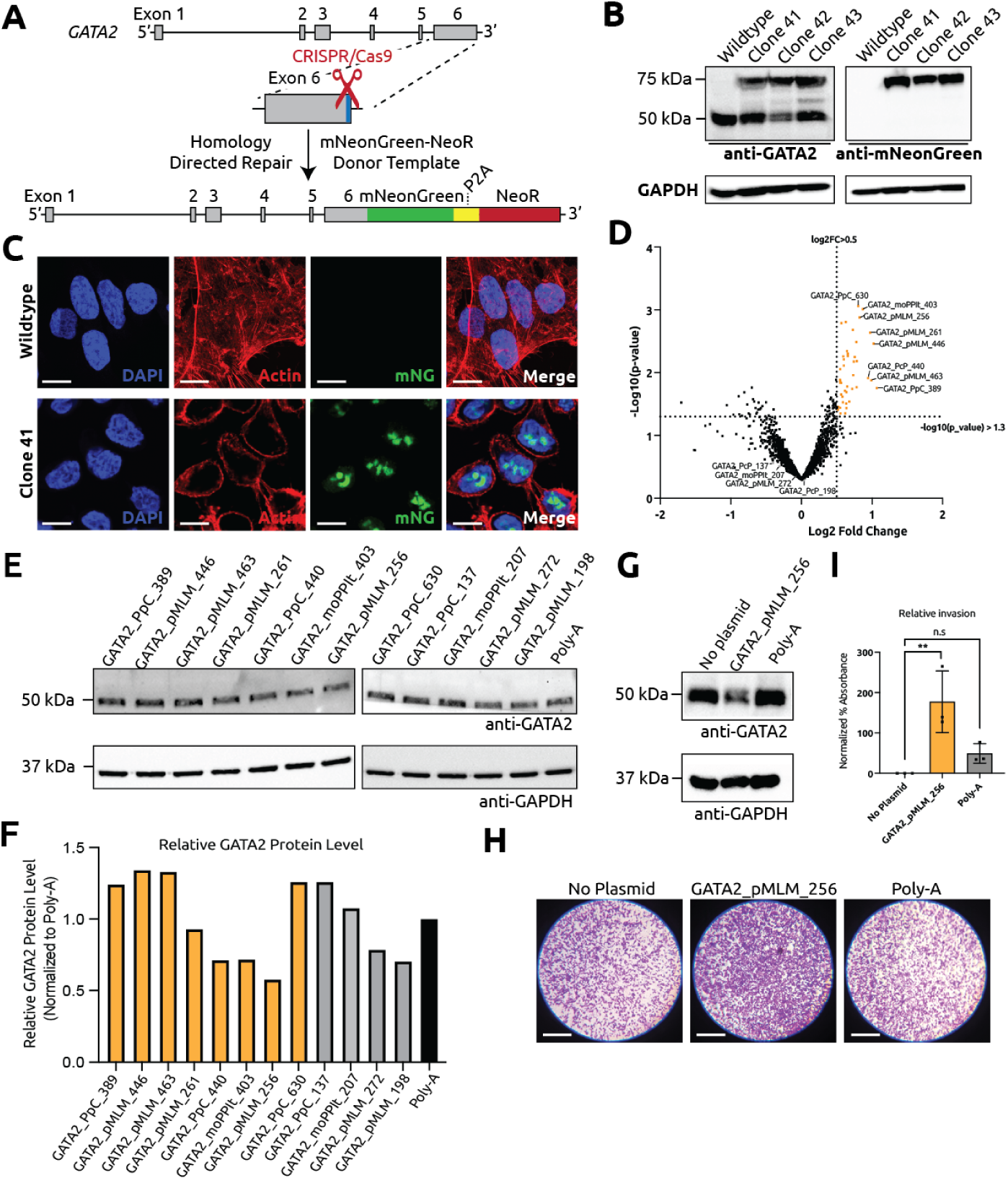
Screening and validation of uAbs against endogenously tagged GATA2. **(A)** CRISPR/Cas9-mediated insertion of mNeonGreen and a neomycin-resistance cassette at the endogenous *GATA2* locus in Ark1 cells. **(B)** Immunoblot validation of GATA2-mNeonGreen expression in wild-type Ark1 cells and three edited clones using anti-GATA2 and anti-mNeonGreen antibodies. GAPDH served as the loading control. **(C)** Immunofluorescence staining of wild-type cells and edited clone 41 showing DAPI, actin, mNeonGreen, and merged channels. Scale bars are shown in the images. **(D)** Volcano plot of peptide enrichment in the GATA2-mNeonGreen screen relative to the mNeonGreen-only control. Yellow points indicate enriched candidates with *p <* 0.05 and log_2_ fold change *>* 0.5. **(E)** Immunoblotting of endogenous GATA2 in Ark1 reporter cells expressing eight enriched candidates, four non-enriched candidates, or polyA. **(F)** Densitometric quantification of GATA2 abundance from **E**, normalized to GAPDH and the polyA control. Yellow bars denote enriched candidates, gray bars denote non-enriched candidates, and the black bar denotes polyA. **(G)** Focused immunoblot validation of GATA2_pMLM_256 relative to no-plasmid and polyA controls. **(H)** Representative crystal-violet staining from the Ark1 cell-invasion assay following expression of GATA2_pMLM_256 or the indicated controls. **(I)** Quantification of the invasion assay in **H**. Bars report mean *±* s.d. from three independent experiments (*n* = 3); n.s., not significant.

We transduced GATA2-mNeonGreen cells and an mNeonGreen-only control line with the doxycycline-inducible GATA2 uAb library. After induction, we collected the lowest 10% of mNeonGreen-expressing cells and compared peptide abundance between the target-tagged and fluorescent-control screens. Eight guides met the GATA2 enrichment criteria of *p <* 0.05 and log_2_ fold change *>* 0.5 (Figure 4D).

We next reconstructed all eight enriched candidates and four non-enriched candidates for secondary validation. Immunoblotting showed that several enriched uAbs reduced endogenous GATA2, with GATA2_pMLM_256 producing the strongest reduction among the enriched candidates tested (Figure 4E,F). Two non-enriched candidates also reduced GATA2 abundance. Thus, the endogenous screen recovered active degraders, although enrichment did not completely separate functional from inactive guides. This partial concordance emphasizes the value of using the screen for candidate prioritization followed by direct validation of endogenous target abundance.

Focused immunoblotting independently confirmed GATA2 reduction by GATA2_pMLM_256 relative to the no-plasmid and polyA controls (Figure 4G). We then advanced this uAb to a cell-invasion assay (Figure 4H). Cells expressing GATA2_pMLM_256 exhibited reduced invasion relative to no-plasmid cells (Figure 4I), supporting further functional evaluation of GATA2-directed degradation in this setting. Together, these experiments validated uAbs against a target expressed under endogenous genomic regulation and thus extend our screening strategy beyond overexpressed target reporters.

## Discussion

The CRISPR-Cas system transformed genome editing by separating target recognition by a guide RNA from the catalytic activity of a Cas effector [40]. This modular architecture has supported gene disruption, single-base editing, gene insertion, and transcriptional regulation [25, 41, 42, 43]. Peptide-guided uAbs apply the same logic to the proteome, with a designed peptide guiding an enzyme effector to the target protein of interest. Similar to high-content CRISPR screening [24], in this work, we coupled this architecture to pooled mammalian selection, enabling functional degraders to be recovered from model-designed peptide libraries in a single screen. We selected four targets to test distinct capabilities of the platform. *β*-catenin established screen performance, endogenous degradation, proteasome dependence, pathway suppression, and degradation kinetics. GFAP and EWS::FLI1 extended the approach to disease-relevant proteins that remain difficult to address through conventional ligand discovery, with EWS::FLI1 providing a stringent test against an extensively disordered fusion oncoprotein. Our GATA2 screen further demonstrated that peptide-guided degraders can be selected against a target expressed from its native genomic locus.

As expected for a functional cellular screen, enrichment varied across targets and did not map uniformly to endogenous degradation. uAb activity depends on peptide binding and expression together with target abundance, localization, and accessibility to E3-mediated ubiquitination, and variation in these factors likely influences recovery from the reporter-low population. The GATA2 results illustrate this relationship. Several enriched guides reduced endogenous GATA2, while two non-enriched candidates also showed activity, supporting the use of enrichment to prioritize candidates for secondary validation (Figure 4D-F). Differences between reporter and disease-model contexts may also influence functional outcomes, particularly for context-dependent fusion oncoproteins such as EWS::FLI1 [44]. We therefore validated selected hits against endogenous proteins and extended the screen to endogenously tagged GATA2, reducing reliance on ectopically expressed target-mCherry reporters that may alter target stoichiometry or epitope accessibility. Focused validation confirmed that GATA2_pMLM_256 reduced endogenous GATA2 relative to the no-plasmid and polyA controls (Figure 4G). GATA2_pMLM_256 did not produce a statistically significant invasion difference relative to polyA (Figure 4H,I), so this experiment most directly establishes the recovery of active degraders against a target expressed under native genomic regulation. Future studies will expand endogenous reporter screening, quantify target engagement and off-target degradation, and incorporate orthogonal functional assays to resolve direct target-dependent effects across cellular contexts.

The modular guide-effector architecture can also extend beyond degradation. We recently developed deubiquibodies (duAbs), which fuse peptide guides to the OTUB1 deubiquitinase domain to stabilize selected proteins by removing ubiquitin conjugates [45]. Together, uAbs and duAbs enable bidirectional control of protein abundance and could support complementary degradation and stabilization screens analogous to CRISPRi and CRISPRa [25]. Our PepRePs architecture extend the same principle to peptide-retargeted phosphatases, enabling guide-directed dephosphorylation [46]. We envision that additional effector domains, including kinases and other post-translational modification enzymes, will broaden the range of programmable protein functions. In parallel, advances in isoform-specific peptide design, conformation-selective guidance, and PTM-aware sequence modeling via methods such as SOAPIA, AlloGen, and PTM-Mamba will further improve control over guide specificity and biological state [15, 16, 47]. Overall our work integrates generative peptide design, functional mammalian selection, and modular enzymatic effectors into a scalable strategy for CRISPR-like proteome perturbation.

## Methods

### Peptide-binder design

Peptide binders were generated using the target sequence-conditioned PepMLM model (https://huggingface.co/ChatterjeeLab/PepMLM-650M) [11], the generative-discriminative PepPrCLIP model (https://huggingface.co/ubiquitx/pepprclip) [12], or the motif-specific moPPIt model (https://huggingface.co/ChatterjeeLab/moPPIt) [14]. All peptide sequences were designed at a length of 20 amino acids.

A previously designed *β*-catenin-targeting peptide, *β*cat_SnP_8, generated using SaLT&PepPr (https://huggingface.co/ubiquitx/saltnpeppr), was included as a positive control [13, 48]. A non-targeting 20-amino-acid poly-alanine sequence (polyA) was used as the standard negative control. All peptide sequences tested in this study are provided in Supplementary Table S1.

### uAb plasmid construction

The uAb plasmid used for validation of enriched peptides was generated from a pcDNA3 vector containing a cytomegalovirus (CMV) promoter and a C-terminal P2A-eGFP cassette. The vector also contained a human phosphoglycerate kinase 1 (hPGK) promoter followed by a blasticidin-resistance cassette. Following PCR amplification with mutagenic primers synthesized by Genewiz, an Esp3I restriction-cloning site was introduced immediately upstream of a flexible GSGSG linker and the CHIPΔTPR coding sequence using KLD Enzyme Mix (NEB).

For the GATA2 screen, the pooled uAb library was constructed with a P2A-mCherry reporter in place of P2A-eGFP. This configuration enabled uAb-expressing cells to be identified by mCherry fluorescence independently of the endogenous GATA2-mNeonGreen reporter.

### Pooled peptide-library design and cloning

For each target, a 2,000-member peptide library was constructed containing polyG and polyA negative controls and model-generated peptide sequences produced using the algorithms described above. The peptide-encoding sequences were synthesized by Twist Bioscience as single-stranded DNA oligonucleotides, amplified by PCR, and cloned into the corresponding uAb architecture using Gibson assembly. The GATA2 library was cloned into the mCherry-containing uAb vector, whereas the *β*-catenin, GFAP, and EWS::FLI1 libraries were cloned into the eGFP-containing uAb vector.

### Lentiviral production

Lentivirus was produced for the target-reporter constructs and pooled uAb libraries. HEK293T cells were seeded in 6-well plates and transfected at approximately 50% confluency. For each well, 0.5 *µ*g of pMD2.G (Addgene #12259), 1.5 *µ*g of psPAX2 (Addgene #12260), and 0.5 *µ*g of the corresponding transfer vector were transfected using Lipofectamine 3000 (Invitrogen) according to the manufacturer’s protocol. The medium was replaced 8 h after transfection, and viral supernatant was harvested at 48 and 72 h after transfection.

### Generation of monoclonal target-mCherry reporter cell lines

For generation of each target-reporter cell line, 1 *×* 10^5^ HEK293T cells were mixed with 20 *µ*L of concentrated lentivirus in a 6-well plate. The medium was replaced 24 h after transduction. Antibiotic selection was initiated 36 h after transduction using 2 *µ*g/mL puromycin (Sigma, P8833). Five days after selection was initiated, the cells were collected for sorting, and individual mCherry-positive cells were deposited into 96-well plates. Following clonal expansion, the genotype of each monoclonal cell line was validated by genomic PCR.

### Generation of endogenously tagged GATA2-mNeonGreen reporter cell lines

Ark1 GATA2-mNeonGreen cells were generated to monitor endogenous GATA2 abundance using an mNeonGreen fluorescent reporter [PMID: 28108553]. Ark1 uterine serous carcinoma cells [PMID: 19920829] stably expressing Cas9 were nucleofected with a guide RNA targeting the stop codon in exon 6 of human *GATA2* and a donor construct encoding mNeonGreen and a neomycin-resistance cassette flanked by approximately 1-kb homology arms.

Following nucleofection, cells were selected with G418 for 2 weeks and sorted according to mNeon-Green fluorescence to isolate individual clones. Clones with the highest mNeonGreen signal were selected for further validation. Monoallelic knock-in at the predicted site within the GATA2 C terminus was confirmed by sequencing. Immunoblotting with anti-GATA2 and anti-mNeonGreen antibodies detected wild-type, untagged GATA2 and a higher-molecular-weight GATA2 species consistent with the predicted size of the GATA2-mNeonGreen fusion protein. The higher-molecular-weight species corresponded to the signal detected with the anti-mNeonGreen antibody. Nuclear localization of GATA2-mNeonGreen was confirmed by fluorescence microscopy.

The Ark1 mNeonGreen-only control cell line was generated by lentiviral transduction with a construct constitutively expressing mNeonGreen and a neomycin-resistance cassette. Following G418 selection, individual clones were isolated, and mNeonGreen expression was confirmed by flow cytometry and fluorescence microscopy.

### Fluorescence microscopy of GATA2-mNeonGreen reporter cells

Cells grown on coverslips were rinsed once with prewarmed 1*×* PHEM buffer and fixed with 4% paraformaldehyde in 1*×* PHEM for 5 min at room temperature. Cells were washed three times with PBS, permeabilized with 0.1% Triton X-100 in PBS for 5 min, and washed twice with 1*×* PHEM. F-actin was stained with Alexa Fluor^TM^ 647-conjugated phalloidin in 1*×* PHEM for 20-30 min at room temperature. Following three washes with 1*×* PHEM, nuclei were stained with DAPI at 1 *µ*g/mL for 5 min. Coverslips were washed, mounted cell-side down using antifade mounting medium, and stored at 4 *^◦^*C protected from light until imaging. Images were acquired using a Nikon inverted microscope equipped with a Yokogawa CSU-W1 spinning-disk confocal system and a CMOS camera.

### Pooled lentiviral-library titration

The titer of each pooled lentiviral peptide library was determined by transducing 6 *×* 10^4^ cells with serial dilutions of lentivirus. Reporter expression was measured 4 days after transduction using an Accuri C6 flow cytometer (BD). GFP fluorescence was measured for the *β*-catenin, GFAP, and EWS::FLI1 uAb libraries, and mCherry fluorescence was measured for the GATA2 uAb library. Each lentiviral library was titrated in the corresponding target-reporter cell line.

### Functional screening of peptide-guided uAbs

Each target-specific screen was performed in duplicate using independent transductions. uAb expression was induced by adding doxycycline at a final concentration of 100 ng/mL. Cells were collected for sorting 5 days after doxycycline treatment. For each replicate, 1 *×* 10^7^ target-mCherry reporter cells were washed once with 1*×* PBS, dissociated using Accutase, passed through a 30-*µ*m CellTrics filter, and resuspended in FACS buffer consisting of 0.5% BSA (Sigma, A7906) and 2 mM EDTA (Sigma, E7889) in PBS. The populations with the highest and lowest 10% of mCherry fluorescence were isolated using an SH800 FACS Cell Sorter (Sony Biotechnology). Genomic DNA was extracted from the sorted cells using the Monarch Genomic DNA Purification Kit (NEB, T3010).

For the GATA2 screen, transduced Ark1 GATA2-mNeonGreen and Ark1 mNeonGreen-only control cells were dissociated into single-cell suspensions using trypsin after 72 h of doxycycline treatment. The cells were briefly incubated in PBS containing DAPI. Approximately 1 *×* 10^6^ cells were sorted from each group. After gating for single, DAPI-negative, mCherry-positive cells expressing the GATA2 uAb library, the 10% of cells with the lowest mNeonGreen fluorescence were collected.

### Peptide-library sequencing

Peptide-encoding sequences were amplified from each genomic DNA sample in 100-*µ*L PCR reactions using Q5 Hot Start Polymerase (NEB, M0493), with 1 *µ*g of genomic DNA added to each reaction. PCR amplification was performed according to the manufacturer’s instructions for 25 cycles using an annealing temperature of 60 *^◦^*C. The primers contained partial Illumina^®^ adapter sequences and were as follows:

Forward: 5*^′^*-ACACTCTTTCCCTACACGACGCTCTTCCGATCTATTGGCTAGCGGATCCGCCACCATG-3*^′^*

Reverse: 5*^′^*-GACTGGAGTTCAGACGTGTGCTCTTCCGATCTAGTTCAGCCGGCCAGAACCGCTGCC-3*^′^*

The amplified libraries were purified using the QIAquick Gel Extraction Kit, and the purified amplicons were submitted to Genewiz for next-generation sequencing.

### Analysis of next-generation sequencing data

Raw sequencing reads from each pooled peptide screen were processed using MAGeCK (Model-based Analysis of Genome-wide CRISPR-Cas9 Knockout; version 0.5.9.5) [34, 35]. Read counts for each peptide were generated using the `MAGeCK count` command. Sequencing reads were mapped to the corresponding uAb library, and counts were normalized across samples.

Differential enrichment was analyzed using the `MAGeCK test` command and the Robust Rank Aggregation (RRA) algorithm. Peptides were ranked according to their fold changes between experimental and control populations, and the rankings were aggregated to identify enriched or depleted sequences. Peptide-level enrichment scores were calculated for negative selection, corresponding to peptide depletion, and positive selection, corresponding to peptide enrichment.

Statistical significance was determined from RRA-derived *p*-values. Multiple-testing correction was performed using the Benjamini-Hochberg procedure to control the false discovery rate. The magnitude and direction of peptide enrichment were reported as log_2_ fold change (log_2_ FC).

### Data visualization

Volcano plots were generated by plotting log_2_ FC against *−* log_10_(*p*) for each peptide. For the *β*-catenin, GFAP, and EWS::FLI1 screens, peptides with log_2_ FC *>* 1 and *p <* 0.01 were classified as enriched. Enriched peptides were shown in red for *β*-catenin, cyan for GFAP, and orange for EWS::FLI1. For the GATA2 screen, peptides with log_2_ FC *>* 0.5 and *p <* 0.05 were classified as enriched and shown in yellow. Peptides that did not meet the corresponding thresholds were shown in gray.

### Cell lysis and immunoblotting

Cells were harvested, detached using 0.05% trypsin-EDTA, and washed twice with ice-cold PBS. Cell pellets were lysed in RIPA buffer (Thermo Fisher Scientific, Cat. #89900) supplemented with protease inhibitor cocktail (MilliporeSigma, Cat. #P8340; 1:100). Lysates were incubated at 4 *^◦^*C for 30 min and centrifuged at 15,000 rpm for 10 min. Protein concentrations were determined using the Pierce BCA Protein Assay Kit (Thermo Fisher Scientific, Cat. #23227).

Twenty micrograms of protein were combined with 4*×* LDS sample buffer containing 5% *β*-mercaptoethanol at a 3:1 ratio and denatured at 95 *^◦^*C for 10 min. Proteins were separated on Bolt^TM^ Bis-Tris Plus gels (Thermo Fisher Scientific, Cat. #NW04125BOX) and transferred using iBlot^TM^ 2 Transfer Stacks. Membranes were blocked in 5% BSA in TBST and incubated overnight at 4 *^◦^*C with rabbit anti-*β*-catenin (D10A8; Cell Signaling Technology, Cat. #8480; 1:1,000), mouse anti-GFAP (SMI 21; BioLegend, Cat. #837201; 1:1,000), rabbit anti-FLI1 (Abcam, Cat. #ab153909; 1:1,000), anti-GATA2 polyclonal antibody [PMID: 16286657], rabbit anti-mNeonGreen (Thermo Fisher Scientific, Cat. #29523-1-AP; 1:8,000), or mouse anti-GAPDH (Santa Cruz Biotechnology, Cat. #sc-47724; 1:10,000).

Primary antibodies were detected using IRDye 800CW donkey anti-rabbit IgG secondary antibody (LI-COR, Cat. #926-32213; 1:2,000) or cross-adsorbed anti-mouse IgG (H+L) Alexa Fluor^TM^ 488 secondary antibody (Invitrogen, Cat. #A-11001; 1:1,000). Membranes were imaged using an Odyssey M imaging system (LI-COR Biosciences, Lincoln, NE, USA). Band intensities were quantified using the gel-analysis function in FIJI, normalized to GAPDH, and expressed relative to the corresponding polyA uAb control.

### Quantitative reverse-transcription PCR

Total RNA was isolated using the Monarch Total RNA Miniprep Kit (NEB, T2010). One microgram of RNA was reverse-transcribed using PrimeScript^TM^ RT Master Mix (Takara Bio, RR036A). Quantitative real-time PCR was performed using SYBR Green detection reagents (Bio-Rad, Richmond, CA) according to the manufacturer’s instructions.

The following primers were used:

EWSAT1 forward: 5*^′^*-TGTGCATTCCGCATCCAGGTGT-3*^′^* EWSAT1 reverse: 5*^′^*-GCTTCAGCAGAGATGTTGCAGG-3*^′^* ACTINB forward: 5*^′^*-CACCATTGGCAATGAGCGGTTC-3*^′^* ACTINB reverse: 5*^′^*-AGGTCTTTGCGGATGTCCACGT-3*^′^*

The amplification program consisted of enzyme activation at 95 *^◦^*C for 10 min, followed by 40 cycles of 95 *^◦^*C for 30 s, 61 *^◦^*C for 30 s, and 72 *^◦^*C for 30 s. Fluorescence was acquired using a QuantStudio 3 Real-Time PCR System (Thermo Fisher Scientific), and EWSAT1 expression was normalized to ACTINB.

### TOPFlash luciferase assay

DLD1 cells were seeded in 96-well plates at a density of 1 *×* 10^4^ cells per well and incubated for 20-24 h before transfection. Each well was transfected with 100 ng of total plasmid DNA using a 1:3 ratio of TOPFlash luciferase reporter plasmid to uAb plasmid. Plasmid DNA was mixed with Lipofectamine 2000 (Invitrogen) in serum-free Opti-MEM (Gibco), incubated for 20 min at room temperature, and added dropwise to each well. After 72 h, luciferase activity was measured using the Luciferase^®^ Reporter Assay System (Promega) according to the manufacturer’s protocol.

### Cell Viability Assay

U251 cells were seeded in opaque 96-well plates at a density of 1 *×* 10^4^ cells per well and incubated for 20–24 h before transfection. Cells were transfected with 100 ng of uAb plasmid DNA per well using Lipofectamine 2000 (Invitrogen) in serum-free Opti-MEM (Gibco). The transfection mixture was incubated for 20 min at room temperature and then added dropwise to the cells. At 72 h post-transfection, 10 *µ*L of WST-8 solution from the Cell Counting Kit-8 (CCK-8; Abcam, catalog no. ab228554) was added directly to each well. The plate was protected from light and incubated for 2.5 h at 37*^◦^*C before absorbance was measured at 460 nm. All measurements were performed in triplicate and normalized to the polyG control.

### Cell-apoptosis assay

TC71 cells were seeded in opaque 96-well plates at a density of 1 *×* 10^4^ cells per well and incubated for 20-24 h before transfection. Each well was transfected with 100 ng of uAb plasmid DNA using Lipofectamine 2000 (Invitrogen) in serum-free Opti-MEM (Gibco). The transfection mixture was incubated for 20 min at room temperature and added dropwise to each well.

After 72 h, 100 *µ*L of 2*×* detection reagent from the RealTime-Glo^TM^ Annexin V Apoptosis Assay (Promega) was added directly to each well, and the plate was protected from light. Apoptosis was monitored using the RealTime-Glo^TM^ Annexin V Apoptosis Assay according to the manufacturer’s instructions. All measurements were performed in triplicate and normalized to polyG control wells.

### Matrigel invasion assay

GATA2 was previously shown to suppress uterine serous carcinoma invasion *in vitro*. For Matrigel invasion assays, 9 *×* 10^4^ cells were transferred to Matrigel-coated membrane inserts with an 8.0-*µ*m pore size (Fisher Scientific, Cat. #8774122) 24 h after plasmid nucleofection. Cells were allowed to migrate for 24 h. The Matrigel was then removed, and the bottom surface of each insert was stained with 0.2% crystal violet prepared in buffered formalin.

Crystal violet was solubilized from the stained Matrigel-coated inserts using a 1:1 mixture of water and methanol. Absorbance was measured at 570 nm, and invasion values were normalized to the no-plasmid control.

## Data Availability Statement

All data needed to evaluate the conclusions of this paper are provided in the main text and supplementary tables. Raw and processed data underlying the graphical figures, including raw immunoblots and sequencing files are provided as Source Data and are available in the following Zenodo repository: 10.5281/zenodo.22087193.

## Author Contributions

L.Z. designed and performed experimental testing, including construct generation, lentiviral monoclonal cell line generation, uAb construct transfections, Western blotting, TOPFlash luciferase assays, qRT-PCR, and cell apoptosis assays, with assistance from L.H. T.C. and S.V. designed guide peptide libraries. D.R. and S.S. generated additional cell lines and performed Western blotting, with supervision from A.V. A.P. performed NGS data analysis. A.M. and D.M. performed GATA2 cell line generation, screening, and analysis. L.Z. and P.C. wrote the manuscript, with input from all authors. P.C. conceived, designed, directed, and supervised the study.

## Supplementary Information

**Figure S1:**
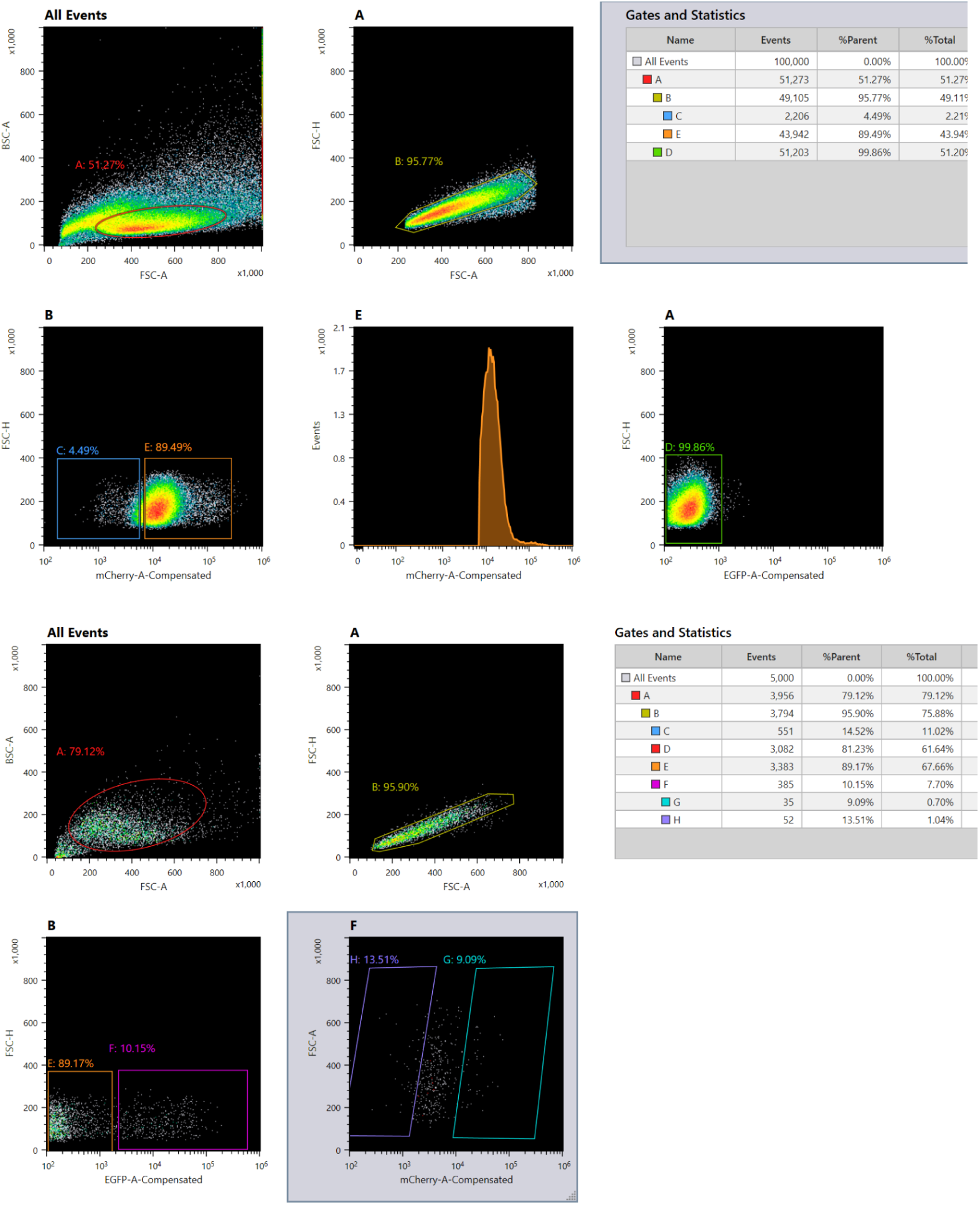
Flow-cytometry gating strategy for pooled uAb screens. We performed spectral compensation using single-color controls before gating and sorting. We first selected the primary cell population and singlets, then gated reporter-positive cells and collected cells within the lowest 10% of target-reporter fluorescence. We sorted approximately 0.5-1.0 million cells for genomic DNA extraction and next-generation sequencing.

**Figure S2:**
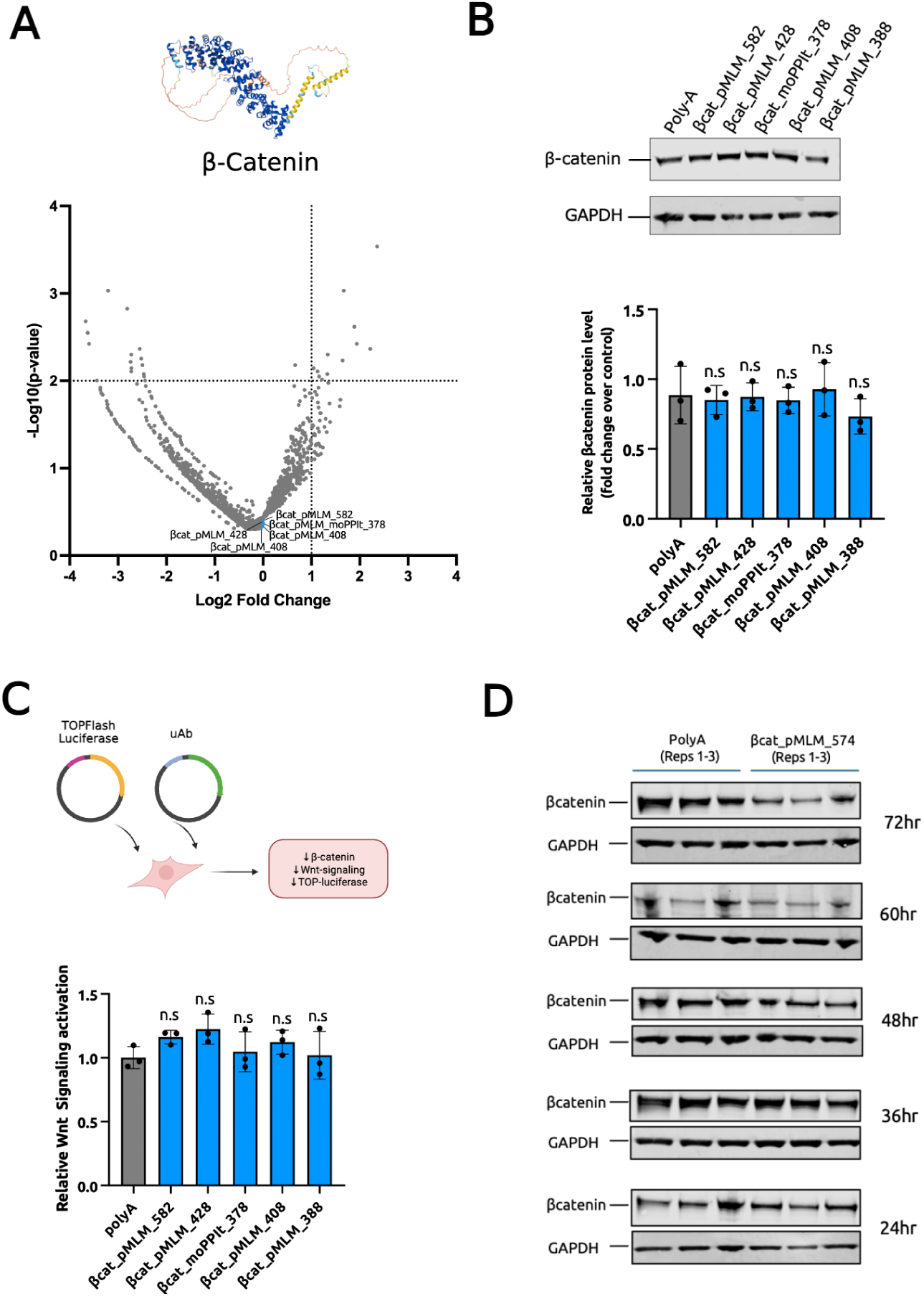
Evaluation of non-enriched peptides for *β*-catenin. **(A)** Volcano plot of peptide enrichment in the *β*-catenin-mCherry reporter cell line. Blue points identify the five non-enriched peptides selected for secondary testing. **(B)** Immunoblotting and densitometric quantification of endogenous *β*-catenin in DLD1 cells expressing *β*cat_pMLM_582, *β*cat_pMLM_428, *β*cat_moPPIt_378, *β*cat_pMLM_408, or *β*cat_pMLM_388. None of the five non-enriched candidates reduced *β*-catenin abundance relative to the polyA control. Bars report mean *±* s.d. from three independent experiments (*n* = 3). **(C)** TOPFlash luciferase assay measuring Wnt-dependent transcription in DLD1 cells. None of the five non-enriched candidates significantly changed Wnt-dependent luciferase activity relative to the polyA control. Bars report mean *±* s.d. from three independent experiments (*n* = 3). **(D)** Time-course immunoblotting of endogenous *β*-catenin in DLD1 cells expressing *β*cat_pMLM_574 or the polyA control. Cells were collected at 24, 36, 48, 60, and 72 h after transfection.

**Figure S3:**
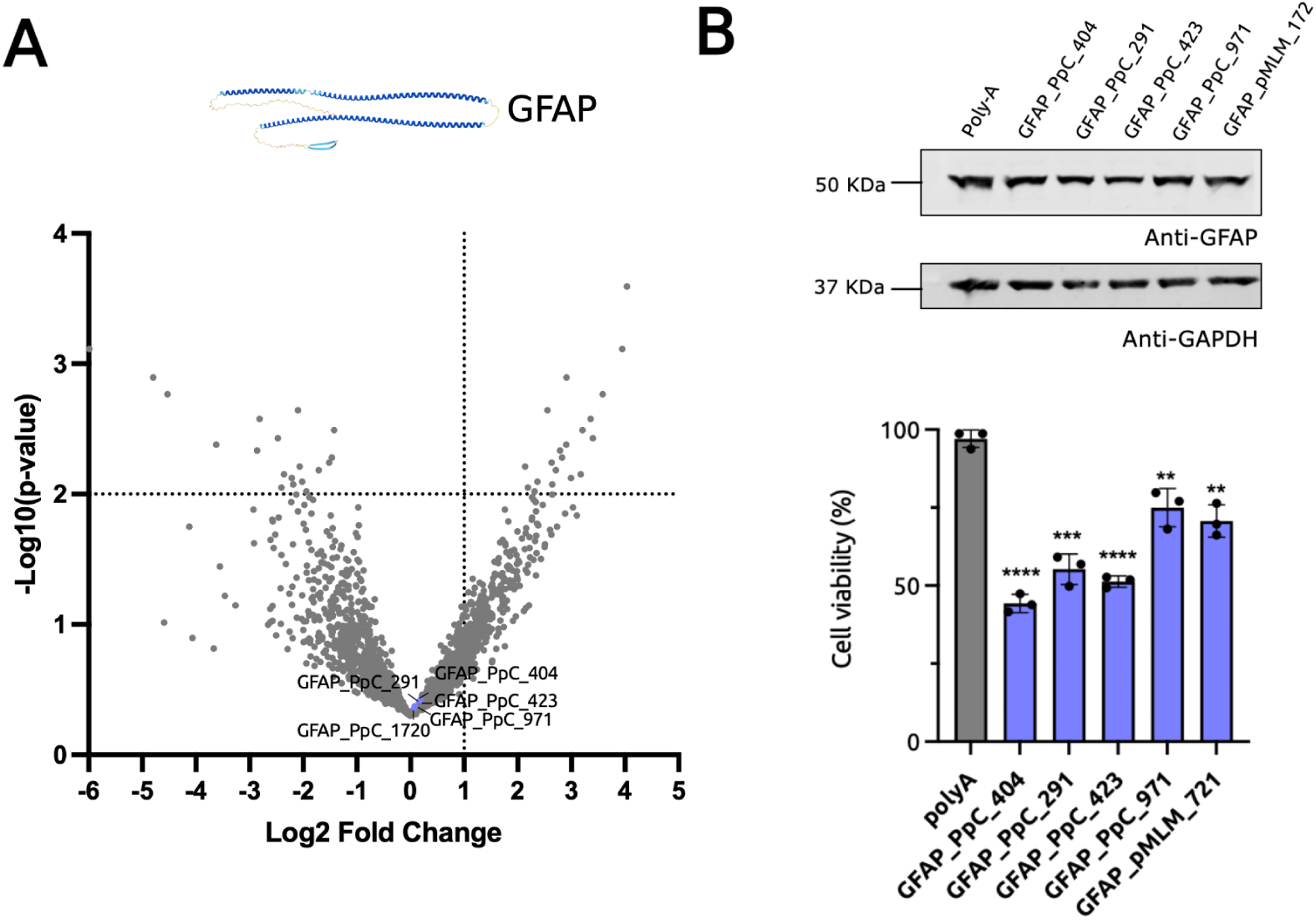
Evaluation of non-enriched peptides for GFAP. **(A)** Volcano plot of peptide enrichment in the GFAP-mCherry reporter cell line. Purple points identify the five non-enriched peptides selected for secondary testing. **(B)** Immunoblotting and densitometric quantification of endogenous GFAP in U251 cells expressing GFAP_PpC_404, GFAP_PpC_291, GFAP_PpC_423, GFAP_PpC_971, or GFAP_pMLM_721. None of the five non-enriched candidates reduced GFAP abundance relative to the polyA control. Bars report mean *±* s.d. from three independent experiments (*n* = 3).

**Table S1:**
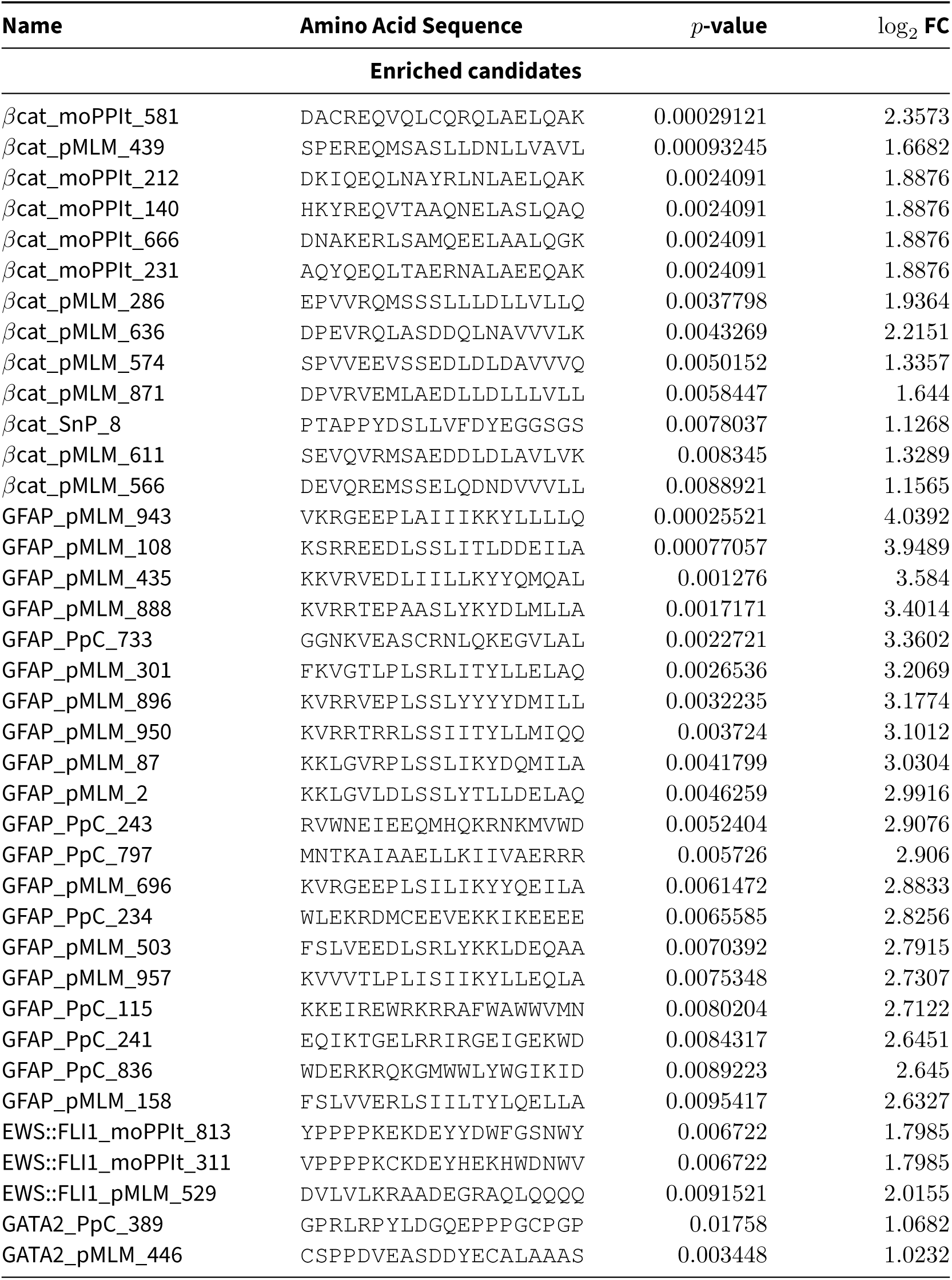
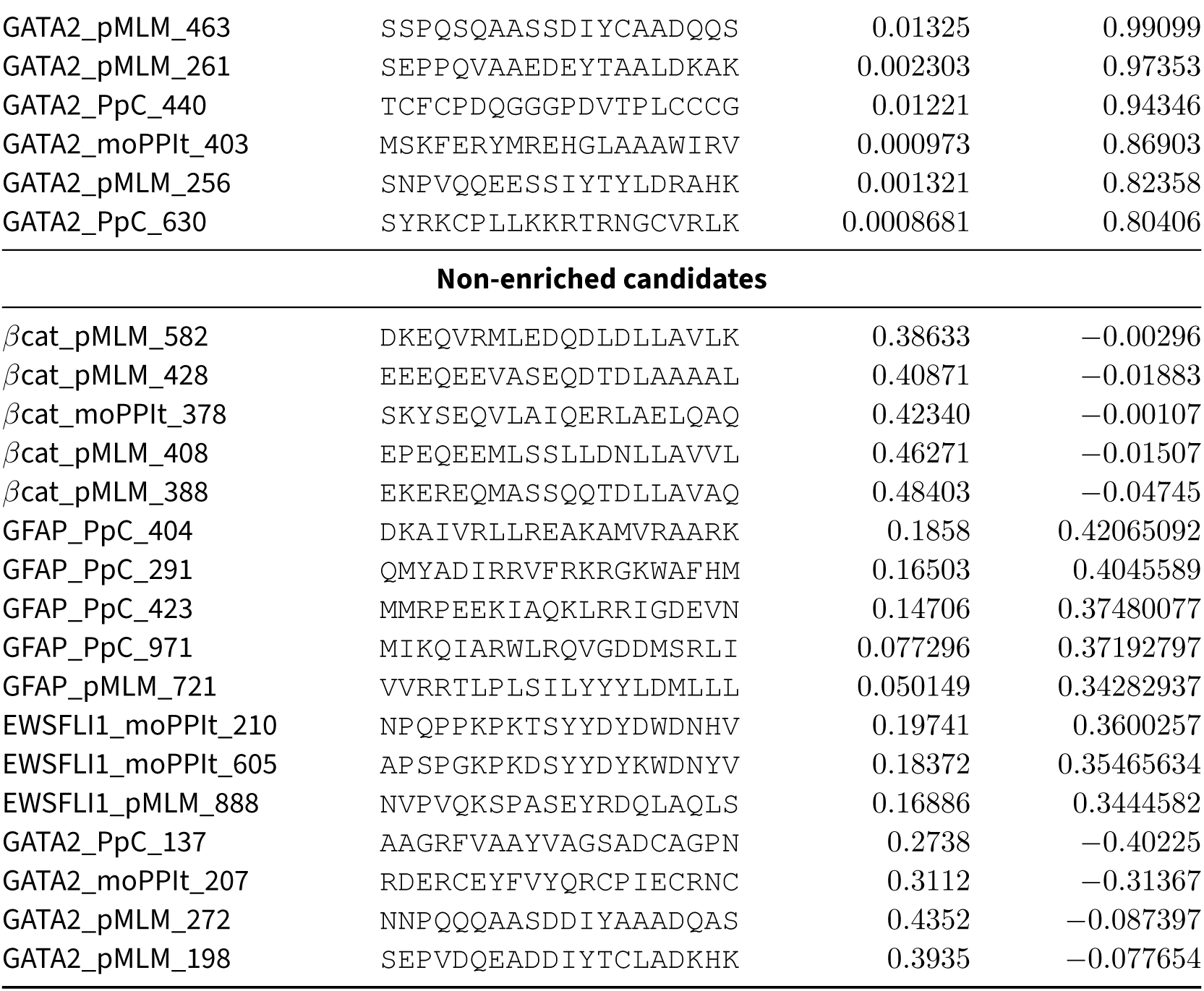
Tested peptide sequences and associated enrichment statistics. The table reports the amino acid sequence, MAGeCK *p*-value, and log_2_ fold change for enriched candidates and the non-enriched candidates used as negative controls.

